# Revealing the co-existence of written and spoken language coding neural populations in the left-ventral occipitotemporal cortex

**DOI:** 10.1101/2024.09.03.610932

**Authors:** Shuai Wang, Anne-Sophie Dubarry, Valérie Chanoine, Julien Sein, Jean-Luc Anton, Bruno Nazarian, Manuel R. Mercier, Agnès Trébuchon, Chotiga Pattamadilok

## Abstract

Reading relies on the ability to map written symbols with speech sounds. The left ventral occipitotemporal cortex (left-vOT) plays a crucial role in this process. Through the automatization of the mapping ability, this specific part of the ventral visual pathway (a.k.a., the Visual Word Form Area) progressively becomes specialized in written word recognition. Yet, despite its key role in reading, the area also responds to speech. This observation raises questions about the actual nature of neural representations encoded in the left-vOT and, therefore, the underlying mechanism of the cross-modal responses. Here, we addressed this issue by applying fine-grained analyses of within- and cross-modal repetition suppression effects (RSEs) and Multi-Voxel Pattern Analyses in fMRI and sEEG experiments. Convergent evidence across analysis methods and protocols showed significant RSEs and successful decoding in both within-modal visual and auditory conditions suggesting that subpopulations of neurons within the left-vOT distinctly encode written and spoken language inputs. This functional organization of neural populations enables the area to respond to speech input directly and indirectly, i.e., after speech sounds are converted to orthographic representations. The finding opens further discussions on how the human brain may be prepared and adapted for an acquisition of a complex ability such as reading.

**Significance Statement:** Learning to read generates new functional responses in neurons in the left ventral visual pathway. Soon after reading acquisition, these neurons become specialized in processing known scripts, thus leading to the functional designation of the “Visual Word Form Area” (VWFA). However, controversies remain regarding the nature of neural representations encoded in this “reading” region, as its activation to speech is also reported. We investigate the neural mechanism(s) underlying these bimodal responses using within and cross-modal repetition suppression and decoding protocols. fMRI and sEEG experiments provided converging evidence indicating that, despite its specialization in reading, VWFA also contained subpopulations of neurons that encode speech. This functional organization could reveal why neurons at this anatomical location are ideal for reading acquisition.

## Introduction

Reading is known to rely on a hierarchical organization of visual information processing: The ventral visual pathway distills information extracted from visual input in successive stages, from early visual cortices that process the physical aspect of the input to the occipitotemporal junction that processes information in a more abstract manner (1–4). Within this process, the left ventral occipitotemporal cortex (left-vOT) is argued to play the central role in recognizing known scripts regardless of their physical characteristics and lesions in this area generally lead to reading deficits (5–12).

The anatomical location of the left-vOT, interfacing the occipital and temporal cortex, as well as its connectivity pattern (13–18) render the area ideal to support the exchanges between visual and non-visual information coming from the left-lateralized spoken language system and other parts of the brain. Several neuroimaging studies indeed showed that in addition to its key role in reading, the area also responds to non-visual language inputs. For instance, studies conducted on congenitally blind readers showed left-vOT responses to Braille script in tactile modality (19) and to letter shapes coded by auditory soundscapes (20). Recent studies conducted in the same population further showed that the left-vOT also responds to non-visual sensory inputs in situations where the inputs could not be translated into spatial or shape patterns, for instance, when blind participants were exposed to vowel sounds (21), spoken words (22, 23) or spoken sentences (22, 24).

In the above examples, such functional reorganization which allows the visual cortex to respond to spoken inputs is mainly attributed to a deprivation of visual sensory input in blinds. Yet, studies conducted on typical populations also showed left-vOT responses to speech in diverse language tasks, ranging from those that required access to spelling knowledge, such as determining whether spoken words contain a target letter or share common rime spellings (25–28), to purely auditory processing tasks such as spoken word recognition (8) and spoken sentence comprehension (29), where an activation of orthographic representations is neither mandatory nor beneficial. These converging findings across levels of language processing, and from both blind and sighted individuals, suggest that some subpopulations of neurons within the left-vOT may be inherently sensitive to speech. Here, we aimed to investigate the underlying mechanism(s) of these cross-modal responses by testing three distinct, but non-mutually exclusive, hypotheses on the nature of representations encoded by neural populations within the left-vOT (Fig. 1A): *Orthographic Tuning hypothesis*, *Multimodal Neurons hypothesis* and *Heterogeneous Neural Populations hypothesis*. As detailed below, although these hypotheses could explain the left-vOT responses to spoken inputs, they make different claims regarding the property of the neurons in the area.

**Fig. 1.**
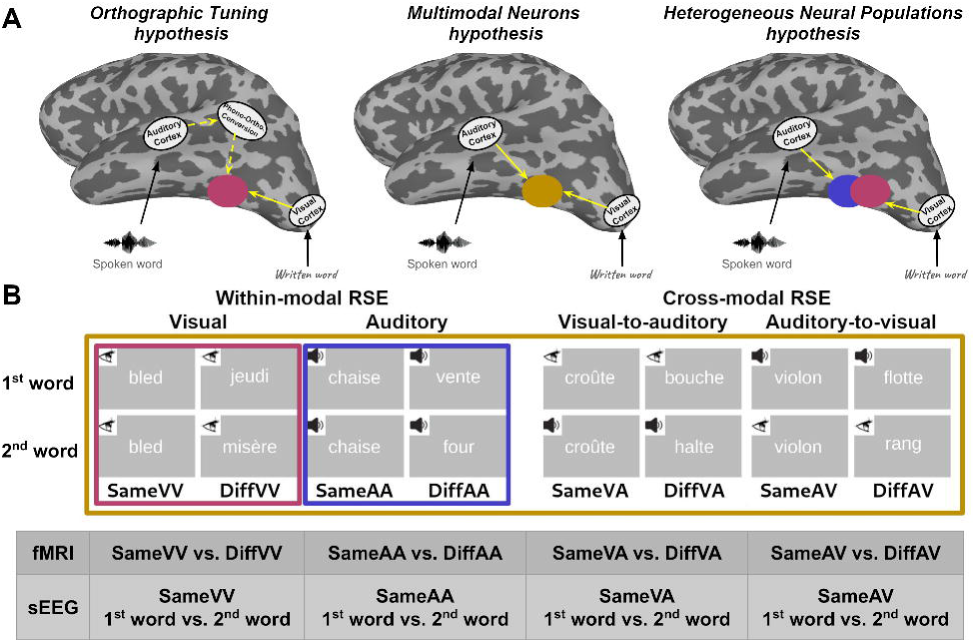
Hypotheses and repetition suppression protocols used in the fMRI and sEEG experiments. (A) Three hypotheses explaining the underlying mechanisms of left-vOT responses to spoken inputs. (B) In the fMRI experiment, eight experimental conditions were used to assess the repetition suppression effects (RSEs). Within-modal visual RSE, within-modal auditory RSE, cross-modal visual-to-auditory RSE and cross-modal auditory-to-visual RSE were assessed using the following contrasts: *SameVV vs. DiffVV, SameAA vs. DiffAA, SameVA vs. DiffVA*, and *SameAV vs. DiffAV*. The sEEG experiment had four experimental conditions containing the same word presented twice in the same (VV, AA) or different modality (AV, VA). Within-modal visual RSE, within-modal auditory RSE, cross-modal visual-to-auditory RSE and cross-modal auditory-to-visual RSE were assessed by contrasting the 1^st^ word vs. 2^nd^ word within the following pairs: *Same VV*, *Same AA, Same VA,* and *Same AV*.

The *Orthographic Tuning hypothesis* (**Fig. 1A** left panel) proposed by Dehaene and colleagues (30, 31) described the left-vOT as a unimodal visual region which, through reading acquisition, became progressively specialized in orthographic coding. This acquisition is assumed to turn neural populations within the left-vOT selectively tuned to known scripts. Although this hypothesis does not predict left-vOT responses to spoken inputs, the authors supplemented a top-down mechanism where spoken inputs could activate orthographic coding neurons in the left-vOT through the conversion of phonological to orthographic representations (8, 32). An alternative view is proposed by the Interactive Account (33, 34) which considers the left-vOT as a convergence area that processes information from both visual regions and spoken language regions. Rather than (or in addition to) responding to orthographic representations generated from spoken inputs, the neural populations within the left-vOT could respond directly to spoken inputs. This account led to two hypotheses: The *Multimodal Neurons hypothesis* (**Fig. 1A** middle panel) assumes that the left-vOT contains multimodal neurons that respond to language inputs regardless of their modality (34). Alternatively, the *Heterogeneous Neural Populations hypothesis* (**Fig. 1A** right panel) assumes that the left-vOT contains different types of unimodal neurons that distinctly encode written and spoken inputs (33).

In a previous study, Pattamadilok and colleagues (35) attempted to disentangle the three hypotheses, using a combination of noninvasive transcranial magnetic stimulation (TMS) and a neural adaptation protocol. Based on the assumption that the behavioral effects of TMS depend on the initial state of the neurons being stimulated (state-dependent TMS effect) (36, 37), the authors reported behavioral evidence (i.e., a facilitation of spoken vs. written word recognition) suggesting that neural populations within the left-vOT, targeted by TMS, were able to adapt their responses to either written or spoken language inputs.

While the above behavioral observation indirectly suggested that the left-vOT might contain distinct populations of neurons that encode either phonological or orthographic information, the present study used functional magnetic resonance imaging (fMRI: Experiment 1) and stereo-electroencephalography (sEEG: Experiment 2) to provide direct neural evidence on the nature of representations encoded in the area. Experiment 1 involved two protocols. First, a repetition suppression protocol aimed to characterize the spatial dimension of within-modal (visual-to-visual; auditory-to-auditory) and cross-modal (visual-to-auditory; auditory-to-visual) repetition suppression effects (RSEs). Based on previous observations that neurons show reduced responses to a repeated presentation of a stimulus or features to which they are sensitive, this protocol has successfully been used to refine the functional resolution of the fMRI signal (38–40). Here, we applied this paradigm to examine the modality of language input encoded by neural populations along the ventral visual pathway, especially within the left-vOT. The three hypotheses described above led to three distinct predictions: 1) The *Orthographic Tuning hypothesis* predicted a reduction of left-vOT activity only in the within-modal visual repetition condition, i.e., where the same written words were repeated, 2) the *Multimodal Neurons hypothesis* predicted a reduction of left-vOT activity in all repetition conditions, regardless of the modality of the inputs, 3) finally, the *Heterogeneous Neural Populations hypothesis* predicted a reduction of left-vOT activity in both within-modal visual and within-modal auditory repetition conditions, but not in the cross-modal conditions. The RSEs were examined by comparing the brain response measured in a pair of different words to the brain response measured in a pair of identical words presented in the same or different modality. The manipulation of word identity and modality resulted in eight conditions depicted in **Fig.1B**. In the second protocol, we conducted a lexical decision task on words and pseudowords presented in either visual or auditory modality. The aim was to complement the repetition suppression protocol by exploring multivariate within- and cross-modal decoding of language inputs according to their lexical category (word vs. pseudoword). In line with the predictions formulated above, the success of the different decoding scenarios (i.e., within-modal visual decoding, within-modal auditory decoding, and cross-modal decoding) would inform us about the nature of the representations encoded in the left-vOT. Experiment 2 used a similar repetition suppression paradigm as Experiment 1 while recording intracranial stereotactic EEG (*sEEG*). The high temporal resolution offered by sEEG signals allowed us to explore the temporal dynamics of the RSEs by comparing the neural time-series on each individual word within the “same word” pairs (cf. **Fig 1B** bottom). Our analyses focused on high-frequency activity (HFA; 70-150Hz), which is strongly correlated with the spiking activity of local neural populations (41,42). A significant reduction of HFA on the second compared to the first word of the pair would indicate an RSE. A combination of these two measures (BOLD and HFA) allowed us to gain insight into the spatial distribution and temporal dynamics of within- and cross-modal RSEs.

## Results

### Experiment 1: fMRI

The fMRI experiment aimed to delineate written and spoken language processing along the ventral pathway covering the left-vOT and to disentangle the three hypotheses on the nature of representations processed by neural populations within this area. To this aim, we first identified the voxels within the left-vOT that responded to written words in a visual localizer task, using a univariate analysis. Then, we further validated previous findings that the same left-vOT voxels also responded to spoken inputs (8, 26–29). Following these steps, we examined the property of the neural populations in the left-vOT and along the ventral visual pathway through the occurrence of within- and cross-modal RSEs. Finally, to examine the neural representations in the left-vOT from a complementary view, we applied multivariate pattern analysis (MVPA) to classify words from pseudowords both in within-modal (visual-to-visual; auditory-to-auditory) and cross-modal (visual-to-auditory; auditory-to-visual) decoding conditions.

### Identification of the regions-of-interest within the left-vOT

Before examining the neural representation within the left-vOT, we first identified, at the group level, the voxels that responded to written inputs, using the *words - consonant strings* contrast from the visual localizer task. The voxel-wise comparison revealed a significant cluster in the left-vOT (FDR q < 0.05; peak MNI - 42, -36, -24; **Fig. 2A**), which was considered in the subsequent analyses as the region-of-interest (henceforth, ROI_GRPvOT_; **Fig. 2B**). In addition to the group-specific ROI identified in the visual localizer task, analyses were conducted on the literature-based ROIs to have a more global view of the neural response along the ventral visual pathway. To this end, we selected 6 ROIs from Vinckier et al. (1) showing a gradient of selectivity from low-level visual features to written words. Those ROIs, located from the anterior to posterior parts of the ventral visual pathway, are referred to as ROI_-40mm_, ROI_-48mm_, ROI_-56mm_, ROI_-64mm_, ROI_-80mm_ and ROI_-96mm_ according to their y coordinates (**Fig. 2B**). Additionally, in the supplementary material (*SI Appendix*, Fig. S1), we presented the analyses conducted on individually defined ROIs (ROI_INDvOT_). These results confirmed those obtained in the ROI_GRPvOT_.

**Fig. 2.**
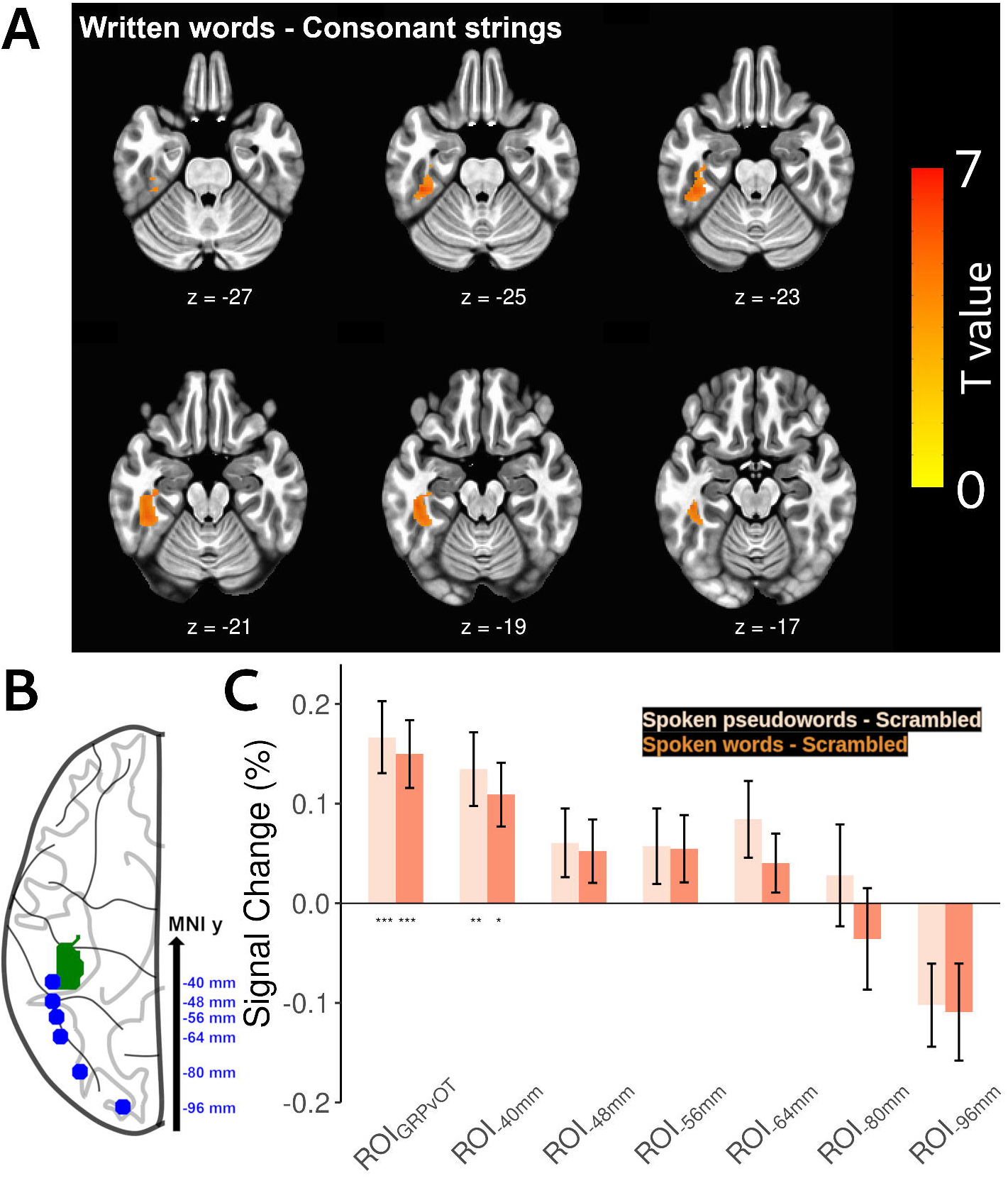
Left-vOT activation in response to written and spoken inputs. (A) Significant activation in the left-vOT revealed by the *written words - consonant strings* contrast from the visual localizer (FDR q < 0.05). (B) Axial view showing the locations of ROI_GRPvOT_ (green patch) and the 6 literature-based ROIs along the ventral pathway (blue dots). (C) Both ROI_GRPvOT_ and ROI_-40mm_ were significantly activated in the *spoken pseudowords - scrambled stimuli* (light orange bars) and *spoken words - scrambled stimuli* contrasts (dark orange bars). This activation induced by spoken inputs progressively decreased from the anterior to the posterior portions of the visual pathway. Error bars represent standard errors. *: *p* < 0.05; **: *p* < 0.01; ***: *p* < 0.005.

### Replication of the left-vOT activation during spoken language processing

Following the ROI identification described above, we examined the left-vOT activation in response to spoken inputs using the *spoken pseudowords - scrambled stimuli*, *spoken words - scrambled stimuli* and *spoken words - spoken pseudowords* contrasts from the auditory task. The ROI-based analysis showed that the ROI_GRPvOT_ and ROI_-40mm_ were significantly activated in both *spoken pseudowords - scrambled stimuli* and *spoken words - scrambled stimuli* contrasts (**Fig. 2C**; *p*s < 0.003 for ROI_GRPvOT_; *p*s < 0.011 for ROI_-40mm_; permutation tests with FWE correction for each ROI) while no significant difference was found in the *spoken words - spoken pseudowords* contrast (all *p*s > 0.058). This observation suggests that the activation was induced by phonological information rather than lexical or semantic one. A similar result was also obtained from the voxel-wise analysis on whole-brain (*SI Appendix*, Fig. S2). Therefore, we successfully replicated previous findings showing that left-vOT voxels that respond to written inputs are also involved in phonological processing of spoken inputs (8, 26–29). Interestingly, as shown in **Fig. 2C**, such involvement decreased progressively from anterior to posterior portions of the ventral visual pathway.

### Investigating the underlying mechanism(s) of left-vOT responses to spoken inputs using within- and cross-modal repetition suppression effects

In the above analysis, we successfully replicated existing observations of the left-vOT response to speech sounds. Here, we attempted to tease apart the three hypotheses on the underlying mechanism(s) of such activation by examining the within- and cross-modal RSEs in the ROIs defined above.

We conducted the ROI-based analyses using permutation tests with FWE correction for each ROI. The result of ROI_GRPvOT_ (**Fig. 3A**) showed significant within-modal RSEs in both visual (*DiffVV* > *SameVV*, *p* < 0.0033) and auditory modalities (*DiffAA* > *SameAA*, *p* < 0.042). However, no significant cross-modal RSE was observed (all *p*s > 0.065). Following this observation, we further explored, at the voxel level, the distribution of clusters of voxels that showed a preference for either within-modal auditory RSE or within-modal visual RSE in the ROI_GRPvOT_ by using the winner-takes-all approach (43, 44). Precisely, for each participant, the β- values were computed for within-modal visual RSE (*DiffVV* - *SameVV*) and within-modal auditory RSE (*DiffAA* - *SameAA*) within each voxel. **Fig. 3B** illustrates the clusters of voxels that showed higher β-values for within-modal visual RSE (in pink) and for within-modal auditory RSE (in purple). The analysis showed a high inter-individual variability in voxel-wise distribution of the two types of within-modal RSE.

**Fig. 3.**
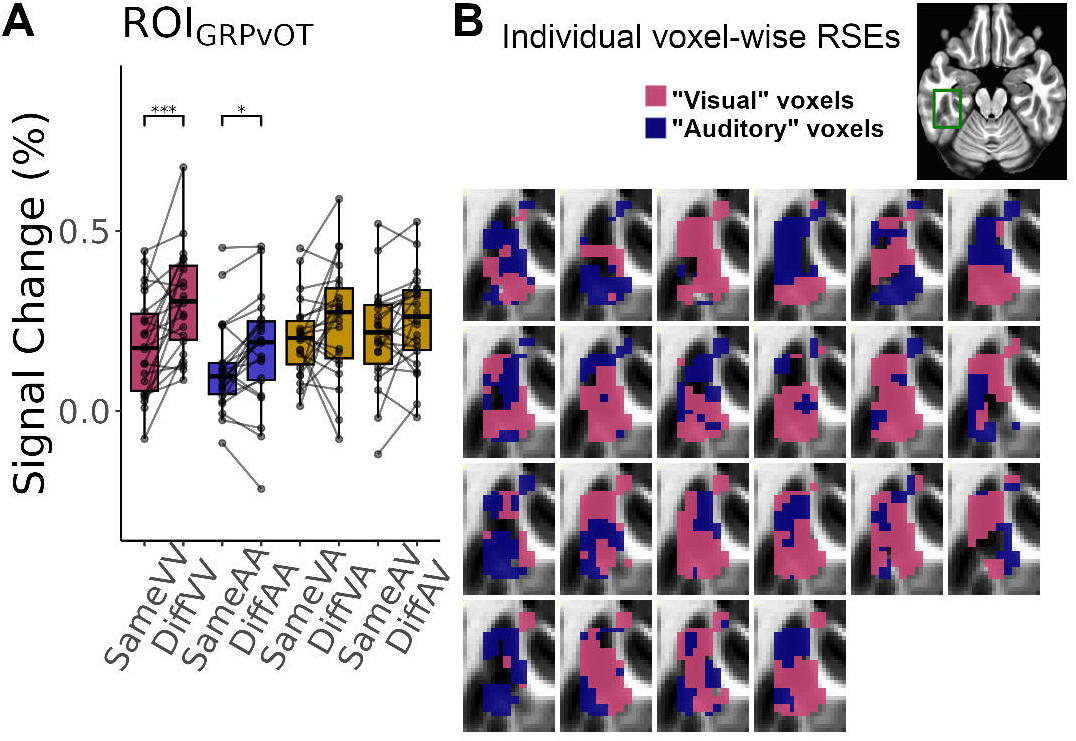
Repetition suppression effects in the left-vOT. (A) Within-modal visual (pink bars) and within-modal auditory RSEs (purple bars) were significant in the ROI_GRPvOT_. No cross-modal RSEs (brown bars) were found. *: *p* < 0.05; **: *p* < 0.01; ***: *p* < 0.005. (B) Within the ROI_GRPvOT_, the winner-takes-all categorization of voxels showed a strong inter-individual variability of the distribution of the voxels with a preference for within-modal visual RSE (“visual” voxels) or within-modal auditory RSE (“auditory” voxels) (MNI y = -21). The spatial location of the depicted results is indicated by the green frame in the axial view at the top-right panel.

The analyses conducted on the 6 literature-based ROIs along the ventral visual pathway showed significant within-modal visual RSEs from ROI_-80mm_ to ROI_-40mm_ (*DiffVV* > *SameVV*: *p* < 0.006, 0.004, 0.009, 0.034 and 0.012, respectively; permutation tests with FWE correction for each ROI). Although no significant within-modal auditory RSE was found in any ROIs (all *p*s > 0.097), their activation profiles showed a trend of an increased RSE for spoken words from the posterior to anterior portions of the pathway (**Fig. 4A**). To further illustrate this trend, the averaged β-values were extracted for the within-modal auditory RSE (*DiffAA - SameAA*) from each slice along the y-axis of a mask of the ventral visual pathway (henceforth VVP mask; **Fig. 4B**). The mask was defined by using the *words - fixation* contrast in the visual localizer with a lenient threshold, *p* < 0.01 uncorrected and then intersected with a pre-defined anatomical mask including the left inferior occipital, inferior temporal, fusiform, lingual and parahippocampal gyri. The result confirmed the gradual increase of RSE for spoken words from posterior to anterior regions (**Fig. 4C**), indicating that the left-vOT might act as a transition area where subpopulations of neurons that respond to spoken or to written language input intermingled.

**Fig. 4.**
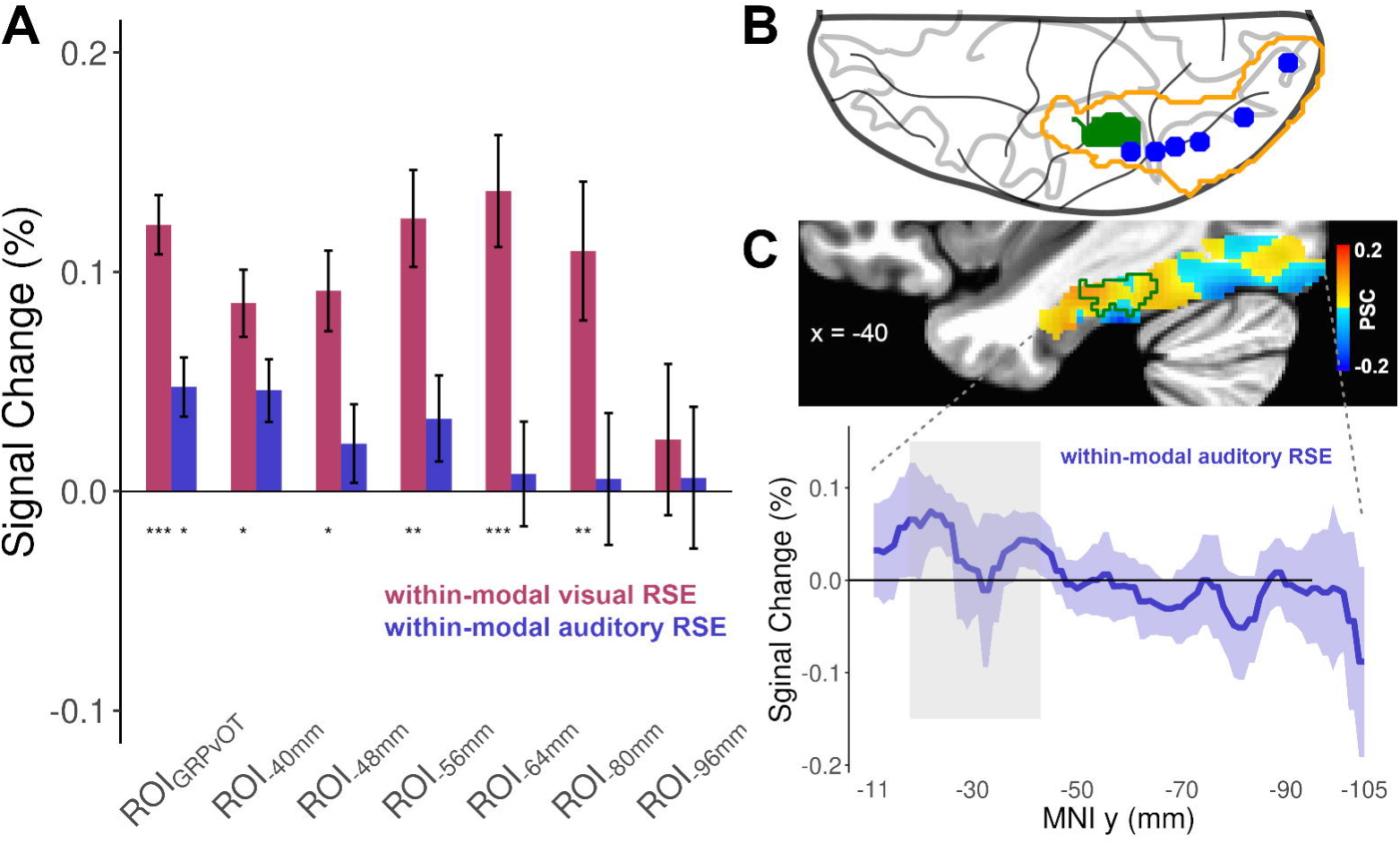
Repetition suppression effects in the ventral visual pathway. (A) The ROI_GRPvOT_ showed significant within-modal visual and within-modal auditory RSEs. The literature-based ROIs from -80mm to -40mm showed significant within-modal visual RSEs, with a trend of a gradual increase of within-modal auditory RSE from the posterior to anterior portions of the ventral visual pathway. (B) Axial view showing the location of the ventral visual pathway mask (orange contour) that contains ROI_GRPvOT_ (green patch) and the 6 literature-based ROIs (blue dots). (C) The brain map illustrates the group-averaged within-modal auditory RSE along the ventral visual pathway (MNI x = -40). The color scale indicates the percent signal change (yellow-to-red indicates positive within-modal auditory RSE). The green contour in the brain map indicates ROI_GRPvOT_. The bottom panel shows the averaged β-values of the within-modal auditory RSE extracted from each slice along the y-axis of the ventral visual pathway mask. Blue ribbon represents standard errors. Gray rectangle indicates the y-axis range of ROI_GRPvOT_.

It is noteworthy that the absence the cross-modal RSE in the analyses conducted on the left-vOT does not call into question the validity of this type of RSE. Indeed, the same analysis conducted on a multimodal language area in left posterior superior temporal sulcus (pSTS) taken from a meta-analysis study on audiovisual integration (45) showed significant RSEs in the four conditions as expected in a multimodal language area. Also as a sanity check, we examined the RSEs in the primary auditory cortex: The area only showed the expected significant within-modal auditory RSEs. These findings which confirmed the validity of the present repetition suppression protocol are reported in the *SI Appendix*, Fig. S3.

### Investigating the underlying mechanism(s) of the left-vOT responses to spoken inputs using within-modal and cross-modal MVPA decoding of stimulus lexicality

The results above suggest the existence of spoken language coding neural populations in the left-vOT by showing a suppression of local neural responses to repeated spoken words. Unlike repetition suppression, MVPA could assess collective responses in neural populations by showing a spatial activation pattern across multiple voxels. As a complementary way to explore the neural representation within the left-vOT, we carried out searchlight MVPA along the ventral visual pathway in a lexical decision task to classify words from pseudowords, based on within-modal (visual-to-visual or auditory-to-auditory) and cross-modal (visual-to-auditory or auditory-to-visual) information. Linear SVM and non-linear SVM classifiers were trained through leave-one-participant-out cross-validation in the following decoding conditions: 1) the classifier was trained and tested on written inputs: visual decoding, 2) it was trained and tested on spoken inputs: auditory decoding, 3) it was trained on written inputs and tested on spoken inputs: visual-to-auditory decoding, and 4) it was trained on spoken inputs and tested on written inputs: auditory-to-visual decoding. According to the *Orthographic Tuning hypothesis*, the left-vOT was expected to show a successful (above-chance-level) classification performance in the within-modal visual condition. The *Heterogeneous Neural Populations hypothesis* predicted a successful decoding in both within-modal visual and auditory conditions. Finally, the *Multimodal Neurons hypothesis* predicted a successful decoding in all conditions.

Within the VVP mask (**Fig 4B**, orange contour), the searchlight analysis using linear SVM only showed two significant clusters with above-chance-level accuracies for written inputs (FWE *p* < 0.05, voxel-wise *p* < 0.005; **Fig. 5A**). The first cluster was centered in the posterior fusiform gyrus extending into the inferior occipital cortex (peak MNI -40, -78, -14). The second one was centered in the anterior fusiform gyrus and largely overlapped with the left-vOT (peak MNI -46, -41, -22). No significant clusters were found for auditory or cross-modal decoding.

**Fig. 5.**
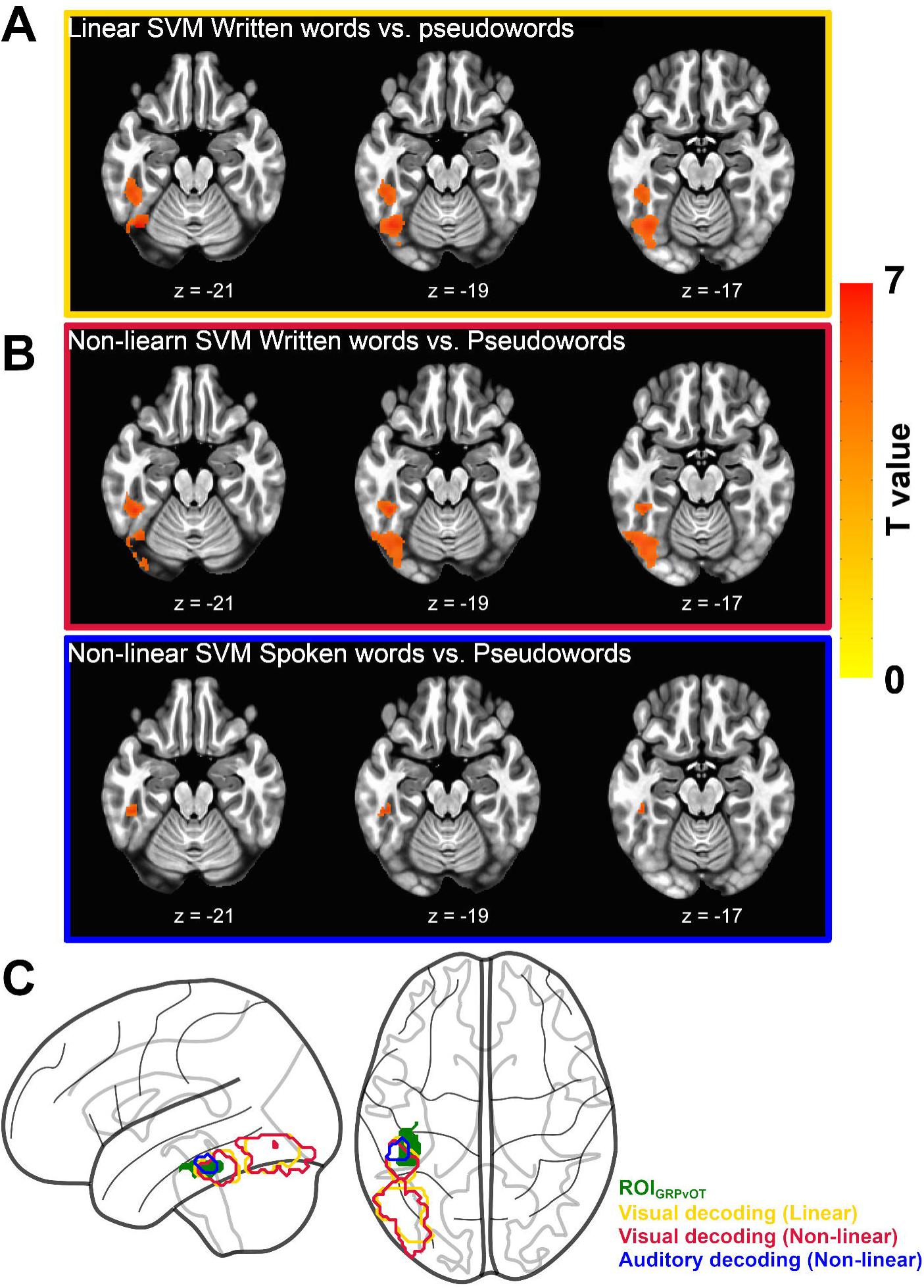
Results of searchlight MVPA for lexicality decoding in the left-vOT (A) Linear SVM revealed two significant clusters with above-chance-level accuracies for written inputs (FWE *p* < 0.05, voxel-wise *p* < 0.005). (B) Non-linear SVM revealed two significant clusters for written inputs and one significant cluster for spoken inputs (FWE *p* < 0.05, voxel-wise *p* < 0.005). (C) Glass brain showing the overlap between the ROI_GRPvOT_ (green patch) and the clusters that showed above-chance visual (yellow and red contour for the linear and non-linear SVM, respectively) and auditory decoding performance (blue contour).

Interestingly, the non-linear SVM revealed a significant cluster that had above-chance-level accuracy for spoken inputs around the left-vOT (**Fig. 5B** and **5C**; FWE *p* < 0.05, voxel-wise *p* < 0.005; peak MNI -51, -34, -19), as well as two clusters with above-chance-level accuracies for decoding written inputs. These two clusters are at the similar locations as those obtained in the linear SVM (FWE *p* < 0.05, voxel-wise *p* < 0.005; peak MNI -54, -64, -9 and -46, -43, -21; **Fig. 5B** and **5C**). Note that neither linear nor non-linear SVM led to an above-chance-level accuracy in the cross-modal conditions. We further applied a simple linear classifier Linear Discriminant Analysis (LDA) and a non-linear Gradient Boosting Classifier (GBC) with searchlight MVPA and confirmed the results obtained above (*SI Appendi*x, Fig S4).

To confirm the validity of the cross-modal MVPAs, the search areas were extended to multimodal language areas (46). The same searchlight analysis was conducted using a mask that covered high-order language regions, i.e., left middle and superior temporal gyri, left supramarginal gyrus and left angular gyrus. The analysis showed a significant decoding performance in all conditions in the left pSTS and left temporoparietal junction (*SI Appendix*, Fig. S5 A and B), i.e., areas reported to be involved in multimodal language processing (47–49). Note that the left pSTS cluster revealed by the searchlight MVPA overlapped with the multimodal ROI reported in Erickson et al.’s meta-analysis on audiovisual integration (45), which also showed significant RSEs (using univariate analyses) in both within-modal and cross-modal conditions (*SI Appendix*, Fig. S3 C and Fig. S4 C).

### Experiment 2: sEEG

#### Exploratory analyses of the temporal dynamics of the within-modal visual and within-modal auditory repetition suppression effects

The converging results obtained in Experiment 1 suggested that the left-vOT responses to written and spoken inputs could reflect the existence of subpopulations of unimodal neurons that are activated by either auditory or visual language modality (35). To complement the above fMRI observations and explore the temporal dynamics of the left-vOT response to spoken and written input, the sEEG recordings of four patients who had electrodes implanted in the left-vOT were analyzed. The modulation of high frequency activity (HFA) within- and cross-modal RSEs was measured as a proxy of population-level local spiking activity (41,42).

To match the location of the areas within the ventral pathway across the fMRI and sEEG experiments, first, we pre-selected the electrodes that were located within a broad box mask covering all individual peaks found in the visual localizer task of the fMRI study (*SI Appendix*, Fig. S1 A). Then, following the same rationale as in the fMRI experiment, we kept the electrodes showing significant HFA to written words. This criterion resulted in a selection of 11 electrodes recorded in the four patients (**Fig. 6A**; see sEEG Electrode Localization and Selection). Then, the 11 electrode signals were analyzed through a multi-patient permutation approach (50, 51). The analysis showed significant within-modal visual and within-modal auditory RSEs [*p* < 0.05, permutation tests with multiple comparisons correction (51)]. The significant within-modal visual RSE was observed in a time-window spanning from 183 ms to 405 ms (duration = 222 ms) and peaked at 206 ms (**Fig. 6B** top panel). The significant within-modal auditory RSE was observed at two distinct temporal stages, i.e., from 219 ms to 254 ms (duration = 35 ms) with a peak at 246 ms, which overlapped with the initial phase of the within-modal visual RSE and from 392 ms to 431 ms (duration = 39 ms) with a peak at 401 ms (**Fig. 6B** bottom panel), which immediately followed the time-window of the within-modal visual RSE.

**Fig. 6.**
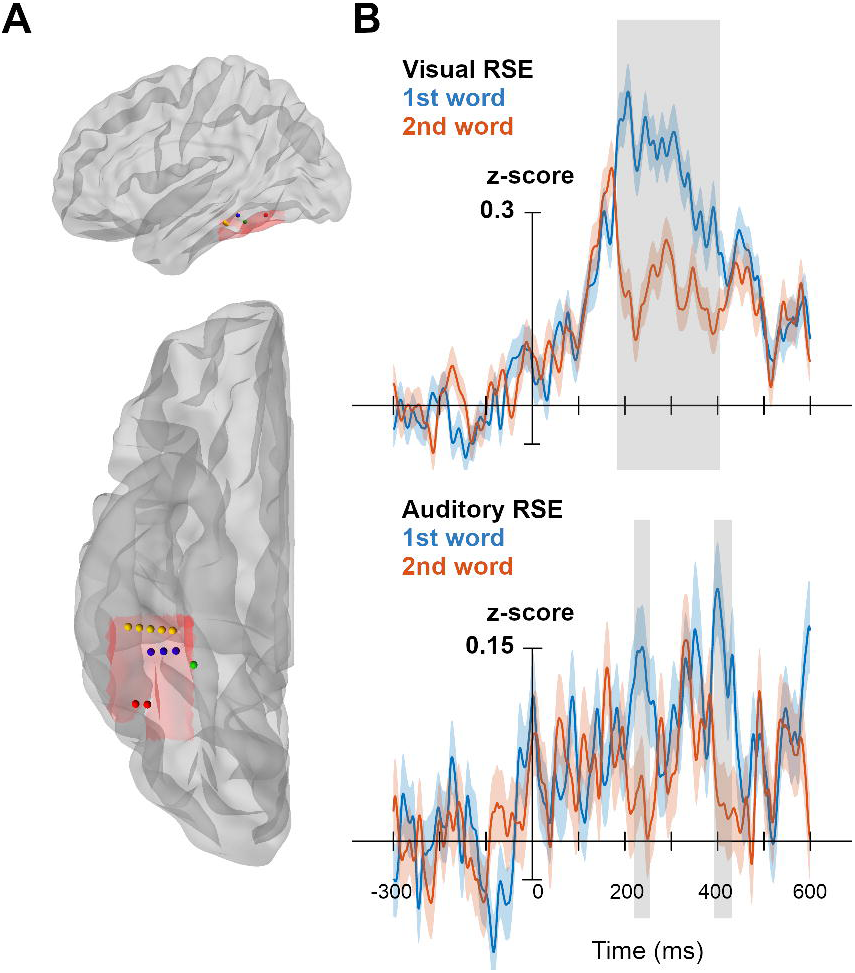
Temporal dynamics of the within-modal visual and within-modal auditory RSEs. (A) The locations of the 11 electrodes in four patients (illustrated in four colors). The red shadow on the cortex view indicates the box mask used for electrode selection. (B) Time courses of high-frequency activity (HFA) recorded on the 1^st^ and the 2^nd^ word of the “same word” pairs presented in the *SameVV* (upper panel) and *SameAA* (lower panel) conditions. The gray bands represent time-windows where the within-modal visual (upper panel) and the within-modal auditory (lower panel) RSEs are significant, i.e., higher brain signal on the 1^st^ than on the 2^nd^ word within a pair [*p* < 0.05, permutation tests with multiple comparisons correction (51)]. Ribbons represent standard errors.

## Discussion

Reading is a bimodal language activity that links written symbols with speech sounds. Mainly underpinned by the left-vOT, a brain structure that interfaces the visual cortex and the temporal language areas, it is not surprising that in addition to its consistent responses to known scripts, this area also responds to speech. The present study examined the underlying mechanism(s) of the left-vOT responses to spoken inputs through fine-grained analyses of both fMRI and sEEG data, aiming to reveal the nature of the representations processed by neural populations within the left-vOT and the temporal dynamics of the process. The analyses presented here provided converging evidence in favor of the *Heterogeneous Neural Populations hypothesis* (33, 34).

Mainly, the repetition suppression protocol in both fMRI and sEEG experiments revealed significant within-modal visual and auditory RSEs, with no reliable evidence for cross-modal RSEs. These two within-modal RSEs respectively suggest the existence of written language coding and spoken language coding neural populations in the left-vOT. In terms of spatial distribution, the winner-take-all analysis conducted on the fMRI data showed that these two types of neural populations intermingled within the area, with a strong inter-individual variability in the distribution of the voxels that showed a preference for either within-modal visual RSE or within-modal auditory RSE. Interestingly, this *Heterogeneous Neural Populations* pattern appears to be specific to the left-vOT, as the other 6 ROIs along the ventral visual pathway only exhibited significant within-modal visual RSEs associated with a trend for an enhancement of within-modal auditory RSE from the posterior to anterior parts of the pathway. This trend is in line with Taylor et al.’s finding (52) that reported a posterior-to-anterior gradient of neural representations reflecting information about phonological form, with the highest level of sensitivity to phonology being observed in the left-vOT.

Regarding the temporal dynamics of the RSEs, the sEEG data revealed a significant within-modal visual RSE over a large time-window and a significant within-modal auditory RSE at two distinct stages that coincided with the onset and the offset of the within-modal visual RSE. Although these results call for a larger scale study, they suggested that the presence of the within-modal auditory RSE in two separated stages might be supported by two distinct mechanisms.

First, the early within-modal auditory RSE that overlapped with the onset of the within-modal visual RSE suggests that the spoken language coding neural populations in the left-vOT may directly receive information from the auditory cortex in the same way as the written language coding neural populations that receive information from the early visual regions. Second, the late within-modal auditory RSE may result from the top-down activation of the written language coding neural populations once the phonological input had been converted into orthographic representations. This second mechanism, which is more time consuming, aligns with the *Orthographic Tuning hypothesis* (8, 32) arguing that the phonological information must be converted into orthographic representations before being processed in the left-vOT.

Nevertheless, the above conclusion also relies on the absence of cross-modal RSE, whose origin remains controversial. As argued in Sawamura et al.’s study (53) in which a single-cell recording was conducted in IT neurons in macaque, “*…for the large majority of the neurons that respond similarly to A and B, the degree of adaptation is smaller for the successive presentation of two different stimuli than for a repetition of a stimulus, even when the two different stimuli (i.e., B and A) elicit, on average, a nearly identical response to their first presentation in a sequence*.” (p. 309). Thus, in the present case, it remains possible that, even in neurons that process both information modalities, the neural adaptation to within-modal repetition, i.e., where the repeated stimuli are physically identical, is still larger than the neural adaptation to cross-modal repetition due to the change in the modality of the sensory inputs. Although no direct measure of within- and cross-modal adaptation at the single-neuron level is available here, we provided pieces of evidence suggesting that the reported absence of cross-modal RSE within the left-vOT could not simply be attributed to the lack of sensitivity of the repetition suppression paradigm. First, we observed a significant cross-modal RSE in the left pSTS which is considered as a multimodal region in language processing (45, 46) (*SI Appendix*). Second, the MVPA result led to the same conclusion as the repetition suppression protocol, i.e., significant within-modal visual and within-modal auditory decoding performances, but non-significant cross-modal decoding performance in the left-vOT. Finally, MVPA also showed a significant cross-modal decoding in the TPJ and pSTS (*SI Appendix*), which indicated that the analysis remained sufficiently powerful to uncover the cross-modal decoding.

From a methodological point of view, repetition suppression and MVPA are two complementary methods (40, 54). While repetition suppression relies on temporally adjunct stimuli and measures neural activity at the level of local neural populations (38–40), MVPA reveals neural representations based on a spatially extended cluster of voxels and, therefore, reflects collective neural responses across voxels within the cluster (55–57). One intriguing observation regarding the MVPA results that deserves further discussion is the difference between linear and non-linear classifiers: When applying MVPA with a linear classifier, only clusters showing successful visual decoding were detected, whereas the non-linear classifier allowed us to uncover clusters in the left-vOT for both visual and auditory decoding. Notably, the visual decoding clusters identified by the linear and non-linear SVM were highly overlapped, thus suggesting that the non-linear classifier not only revealed the same neural representations as did the linear classifier, but also those that the linear classifier failed to detect. The advantage of non-linear over linear classifiers could be due to their flexible decision boundary (58, 59). In contrast to the predominant written language coding neural populations in the left-vOT, the spoken language coding neural populations could be more sparsely distributed. Therefore, a flexible decision boundary offered by the non-linear classifier (e.g., polynomial, sinusoid) might show some additional benefits in capturing such complex combination of activity across voxels.

Overall, we provided converging findings from univariate activation, fMRI/sEEG repetition suppression and non-linear MVPA that support the co-existence of written and spoken language coding neural populations within the left-vOT. These observations are in line with the previously observation from a TMS adaptation paradigm (35). Under the multisensory convergence framework proposed by Meredith (60), the co-existence of the two unimodal neural populations within the left-vOT reflects an “areal convergence”. In other words, written and spoken inputs may not converge to the same (multimodal) neurons, but to the same area where the two unimodal neural populations intermingled. Nevertheless, at this stage of research, one could not rule out the existence of multimodal neurons (neural convergence) (60). Such evidence could be obtained using a technique with higher spatial resolution like single-cell recording.

The converging findings obtained across brain imaging techniques (TMS, fMRI, sEEG) opens further discussions both on the origin of the Visual Word Form Area and on the consequence of reading acquisition on functional reorganization of the human brain. Prior research suggests that the emergence of reading specialization in the left-vOT is triggered by the mapping between the orthographic and phonological codes during reading acquisition (61–63). Yet, it remains to be investigated whether this repeated connection between the two language codes has rendered some “visual neurons” sensitive to speech input. In such a case, the existence of the spoken language coding neural populations in the left-vOT would be a *consequence* of functional reorganization of the visual pathway following reading acquisition (64, 65). An alternative possibility is that these spoken language coding neurons may *predate* reading acquisition. In the other words, some neurons that are located at the transition between the visual and the spoken language system, and that become the Visual Word Form Area later on, might already show some degree of sensitivity to spoken language before reading acquisition. This assumption seems plausible given existing findings on brain connectivity. Indeed, a longitudinal study reported that the anatomical connectivity between the left-vOT and the temporal language areas precedes reading acquisition, and the anatomical connectivity pattern of the left-vOT with the rest of the brain can predict the precise location of the area that will become the Visual Word Form Area once reading is acquired (18). Moreover, the left-vOT shows functional connectivity with spoken language regions in neonates as early as one week after birth (66). Given that cross-modal projections can trigger responses to new sensory modalities even in a unimodal area (67, 68), the connections between the left-vOT and spoken language regions could potentially lead to the development of spoken language coding neural populations in the visual pathway independently of reading acquisition. In either way, these neurons could facilitate the acquisition of orthographic-phonological mapping when one learns to read. This potential benefit is obvious at the initial stage of reading acquisition, as numerous studies have shown a positive relationship between reading skill in children and the activation of the left-vOT during spoken language processing (62, 69–71).

Finally, the co-existence of written and spoken language coding neural populations also offers a new perspective for understanding the connections of the left-vOT with other brain regions (13–15, 72). For instance, in spoken sentence processing, the left-vOT was reported to act as a bridge linking distributed brain regions and to adapt its connectivity pattern to task demands and quality of speech signal (15). This adaptive pattern of connectivity in speech processing tasks could be supported by different mechanisms depending on the types of neurons, i.e., either through a direct communication between the phonological coding neural populations and the spoken language system or through a top-down activation of the orthographic coding neurons once speech sounds have been converted to orthographic representations (8, 64). This perspective suggests a potential benefit of considering the different types of neurons in fine-grained connectivity analyses.

In summary, the left-vOT exhibited significant activation, repetition suppression effects and successful decoding performance to both written and spoken inputs. Our study provided converging evidence for the co-existence of written and spoken language coding neural populations in the left-vOT, in line with the *Heterogeneous Neural Populations hypothesis* (33, 34). These observations not only provide insight into the nature of the representations encoded in the most important area of the reading network, but also open further discussions on how the human brain may be prepared and adapt for an acquisition of a complex ability such as reading.

## Materials and Methods

### Experiment 1: fMRI

#### Participants

Twenty-two native French speakers participated in the study (age mean±SD: 26.0±4.3, 13 females). All participants were healthy, right-handed, with normal hearing and vision and reported no neurological or language disorders. Written informed consents were obtained from all participants. The study was approved by the national ethics committee (CPP Sud-Méditerranée, n° ANSM 2017-A03614-49).

#### Stimuli

Word stimuli were mono- or disyllabic nouns selected from the French database LEXIQUE (http://www.lexique.org) with a minimal lexical frequency of 5 per million. In all tasks, when different subsets of words were used in different experimental conditions, they were matched on number of letters, number of syllables, number of phonemes, lexical frequency, OLD20, PLD20 and uniqueness point. All spoken inputs were recorded by a native French female speaker using an AKG C1000S microphone in an anechoic chamber (Centre d’Expérimentation sur la Parole, Laboratoire Parole et Langage, Aix Marseille University, CNRS), with a sampling rate of 48 kHz (32 bits).

#### Visual localizer task

192 words were selected from the database and 96 consonant strings were created by matching the number of letters with the words. All stimuli were presented in the visual modality.

#### Auditory task

96 words were selected and 96 pseudowords were generated by Wuggy (73), using the same phonemes as in the words. Then, 96 scrambled stimuli were generated from the 96 spoken pseudowords by permuting Fourier components to remove phonological information while preserving acoustic information. All stimuli were presented in the auditory modality.

#### Repetition suppression protocol

144 words were selected as NoGo trials and assigned to eight conditions as illustrated in **Fig. 1B**. From this initial word pull, 72 words were generated in the visual modality and 72 words were generated in the auditory modalities. The aforementioned word properties were matched between the eight conditions (Kruskal-Wallis test; all *p*s > 0.49). Eight pseudowords (four in the visual modality and four in the auditory modality) were generated using Wuggy (73). They were used as Go trials.

#### Lexical decision task (for MVPA)

60 words were selected and 60 pseudowords were generated by Wuggy (73), using the same phonemes as in the words. All stimuli were generated in both visual and spoken modalities.

#### Procedures

For each participant, we conducted a single scanning session in fMRI comprising four tasks: 1) a block-design visual localizer task to localize the voxels within the left-vOT that responded to written words, 2) a block-design auditory task to examine whether the voxels identified in the visual localizer also responded to spoken inputs, 3) an event-related pseudoword detection task in which pseudowords were randomly included among sequences of words which were presented in within- and cross-modal repetition suppression conditions (**Fig. 1B**) and, 4) an event-related lexical decision task using both written and spoken words and pseudowords for MVPA.

#### Visual localizer task

The visual localizer task was a block-design task presented in a single run that lasted 6.60 min. Words and consonant strings were presented in separate blocks of 11.8s each. Each block contained 24 stimuli of the same condition. Each stimulus was displayed for 328 ms, followed by a 164 ms blank screen. Altogether, there were eight blocks of words and eight blocks of consonant strings, interleaved with 16 baseline blocks of “fixation” in which a cross remained on the screen for 11.8s. No conditions were repeated twice in a row. During the task, participants were required to press a response button when they detected the target stimuli (######), which appeared randomly eight times between blocks. Each target stimulus lasted 328 ms and was followed by a fixation (jittered from ∼1.2s to 1.8s) to allow responses. All stimuli were centrally presented on the screen in white font on a gray background.

#### Auditory task

The auditory task was a block-design task presented in a single run that lasted 6.73 min. Spoken words, spoken pseudowords and scrambled stimuli were presented in separate blocks of 12.1s each. Each block contained 12 stimuli of the same condition. Each stimulus was presented for 804 ms, followed by a 201 ms silence. Altogether, there were eight blocks of spoken words, eight blocks of spoken pseudowords, eight blocks of scrambled stimuli and 16 silent-rest baseline blocks (which also lasted 12.1s). All blocks were presented pseudo-randomly to avoid repetition of the same condition. During the task, participants were required to press a response button when they detected the target stimuli (beep sounds), which appeared randomly eight times between blocks. Each target stimulus lasted 335 ms, followed by a fixation (jittered from ∼1.2s to 1.8s) to allow responses. All stimuli were presented through insert earphones.

#### Repetition suppression protocol

The protocol was an event-related design with eight conditions based on word identity (same vs. different) and modality (auditory/auditory, visual/visual, auditory/visual or visual/auditory), as illustrated in **Fig. 1B**. Four types of repetition suppression effect (RSE) were estimated by contrasting the brain activity measured during the processing of same-word pairs with the activity during the processing of different-word pairs : 1) Auditory RSE, all stimuli were presented in the auditory modality (i.e., *SameAA vs. DiffAA*); 2) Visual RSE, all stimuli were presented in the visual modality (i.e., *SameVV vs. Diff VV*); 3) Auditory-to-visual RSE, in both same-word and different-word pairs, the first stimulus was presented in the auditory modality and the second in the visual modality (i.e., *SameAV vs. DiffAV*), and 4) Visual-to-auditory RSE, in both same-word and different-word pairs, the first stimulus was presented in the visual modality and the second in the auditory modality (i.e., *SameVA vs. DiffVA*). The task was presented in two runs that lasted 8.2 min each. Each run contained 9 trials from each condition. The resulting 72 trials were presented pseudo-randomly to avoid repetition of the same condition. Each trial lasted 1886 ms, including a pair of 820 ms words separated by a 246 ms interval. The inter-trial interval was jittered from 7298 ms to 9594 ms. During the task, the participants were not informed about the existence of word pairs and were required to press a button when they detected pseudowords (Go trials), which were randomly presented in either modality.

#### Lexical decision task (for MVPA)

The lexical decision task was an event-related design with four conditions, i.e., written words, written pseudowords, spoken words and spoken pseudowords. The task was presented in five runs that lasted 3.8 min each. Each run contained 12 trials from each condition. The resulting 48 trials were pseudo-randomly presented, with a constraint that each stimulus was presented only once in each run in either visual or auditory modality. Each trial lasted 820 ms with an inter-trial interval jittered from 3034 ms to 4182 ms. During the task, the participants were required to determine whether a stimulus is a word or a pseudoword by pressing the corresponding button.

#### Data Acquisition

The experiment was conducted on a 3T Siemens Prisma Scanner (Siemens, Erlangen, Germany) at the Marseille MRI center (Centre IRM-INT@CERIMED, UMR7289 CNRS & AMU, http://irmf.int.univ-amu.fr/) using a 64-channel head coil. T1-weighted images were acquired using an MPRAGE sequence (voxel size = 1×1×1 mm^3^, data matrix = 256×240×192, TR/TI/TE = 2300/900/2.88 ms, flip angle = 9°, receiver BW=210Hz/pix). Fieldmap images were obtained using Dual echo Gradient-echo acquisition (TR = 677 ms, TE1/TE2 = 4.92/7.38 ms, FOV = 210×210 mm^2^, voxel size = 2.2×2.2×2.5 mm^3^). Whole brain functional images were collected during the auditory task using a gradient EPI sequence (TR = 1206 ms, TE = 30 ms, 54 slices with a thickness of 2.5 mm, FOV = 210×210 mm^2^, matrix = 84×84, flip angle = 65°, multiband factor = 3). Partial coverage functional images were collected during the other tasks to increase the spatial resolution of the ventral occipito-temporal cortex, using a gradient EPI sequence (TR = 1148 ms, TE = 32 ms, 42 slices with a thickness of 1.75 mm, FOV = 210×210 mm^2^, matrix = 114×114, flip angle = 66°, multiband factor = 3). Auditory stimuli were presented to the subject via the Sensimetrics S14 MR-compatible insert earphones with a Yamaha P-2075 power amplifier.

#### Data pre-processing

Pre-processing was conducted by using fMRIPrep 20.2.0 (74). For more details on the preprocessing pipeline, see fMRIPrep’s documentation (https://fmriprep.org/en/20.2.0/workflows.html) and *SI appendix, SI Methods*.

#### Data analysis

Pre-processed functional data were scaled to percent of signal change and modeled using voxel-wise GLM for each task per participant. Motion contaminated volumes were identified and censored along with the prior volume if their FD > 0.5mm. On average, 0.84% of the volumes were censored for the visual localizer task, 0.68% were censored for the auditory task, 0.61% were censored for the word repetition task and 0.37% were censored for the lexical decision task. The six motion parameters, their temporal derivatives, and all their corresponding squared time series (i.e., 24 head motion regressors) were included in the GLMs to control for the impacts of head motion (75). In addition, the first six principal components of white matter and of CSF extracted by the aCompCor method (76) were used as nuisance regressors for the GLMs to reduce influence of physiological noise. The cosine-basis regressors estimated by fMRIPrep for high-pass filtering were also included in the GLMs as nuisance regressors.

#### Regions of interest (ROIs) from the visual localizer task

Functional ROIs of the left-vOT were extracted from T-maps contrasting *words* and *consonant strings*. T-maps were thresholded at both group and individual level. At the group-level, the left-vOT (ROI_GRPvOT_) was defined as ROI by identifying a single cluster of voxels through adjusting the significant threshold from a lenient *p* < 0.05 unc. to FDR q < 0.05 (equivalent *p* < 7.7e-4). At the individual-level, the left-vOT (ROI_INDvOT_) was defined as ROI by creating a sphere of 8mm radius around the peak coordinates obtained from the individual contrast with *p* < 0.001. The mask of the ventral visual pathway was defined by using the *words - fixation* contrast in the visual localizer with a lenient threshold, *p* < 0.01 uncorrected and then intersected with a pre-defined anatomical mask including the left inferior occipital, inferior temporal, fusiform, lingual and parahippocampal gyri in the Automated anatomical labelling atlas (77).

#### Repetition suppression effect

The GLM for the word repetition task included eight regressors of interest, that are *SameAA*, *DiffAA*, *SameVV*, *DiffVV*, *SameAV*, *DiffAV, SameVA* and *DiffVA*. The auditory and visual RSEs were estimated by the contrasts of *<u>DiffAA - SameAA</u>* and *<u>DiffVV - SameVV</u>*, respectively. The cross-modal auditory-to-visual and visual-to-auditory RSEs were estimated by the contrast of *<u>DiffAV - SameAV</u>* and *<u>DiffVA - DiffVA</u>*, respectively.

#### Multi-voxel pattern analysis (MVPA)

In order to conduct MVPA, trial-wise estimates (i.e., β coefficients) were extracted for each condition in the lexical decision task by using the Least Squares — Separate (LSS) method (78) which ran a GLM for each trial (3dLSS in AFNI). Trials with incorrect responses were excluded from the analysis, resulting in 9.7% trials being rejected on average. The MVPA was then conducted using Scikit-learn (79) and Nilearn v0.10.1 (80). Linear and non-linear SVM classifiers, Linear Discriminant Analysis (LDA) and Gradient Boosting Classifier (GBC) in Scikit-learn were trained with default parameters to classify words from pseudowords through leave-one-participant-out cross-validation. The searchlight with a sphere of 4mm radius was applied within the VVP mask defined by the visual localizer. The individual accuracy maps were entered in the group tests to compare to the chance level accuracy (50%).

Unless stated otherwise, the voxel level statistical comparisons in the present study were conducted by using AFNI’s *3dttest++* and *3dClustSim* with FWE cluster-based correction *p* < 0.05, voxelwise *p* < 0.005. The ROI-based statistical comparisons were conducted by using pairwise permutation tests implemented in R and its *coin* (http://coin.r-forge.r-project.org/) and *rcompanion* (http://rcompanion.org/) packages and were FWE corrected per ROI.

### Experiment 2: sEEG

#### Participants

Eleven patients who underwent intracranial EEG monitoring for presurgical evaluation of epilepsy at the Hôpital de La Timone (Marseille, France) were recruited (age mean±SD: 29.8±12.2, 6 females). Four patients with electrodes located within the left-vOT were included in the present study (age mean±SD: 26.3±5.7, 2 females; see sEEG Electrode Localization and Selection). None of the patients had previously undergone brain surgery. No seizures were observed within the 24 hours preceding the experiment. Written informed consents were obtained from all patients. The study was approved by the Institutional Review Board of the French Institute of Health (IRB00003888).

#### Stimuli

Critical word stimuli had the same characteristics as those described in Experiment 1. The spoken inputs were also recorded in the same condition. In this experiment, 200 words were selected and assigned to four conditions of the repetition suppression protocol as illustrated in **Fig. 1B**. Thus, from the initial pool of 200 words that had been selected, 50 words were in visual modality, 50 words were in auditory modality and 100 words were in both modalities. The number of letters, number of syllables, number of phonemes, lexical frequency, OLD20, PLD20 and uniqueness point were matched between the four conditions (Kruskal-Wallis test; all *p*s > 0.40). In addition to the critical words, the material also contained other 400 fillers (50 written words, 50 written pseudowords, 50 consonant strings, 50 spoken words, 50 spoken pseudowords, 50 scrambled audio stimuli, 50 videos of lip-movements with speech sounds and 50 videos of lip-movements without speech sounds). These stimuli were analyzed in other studies and were included here to reduce participants’ expectation on stimulus repetition.

#### Procedures

The RSE protocol applied in this experiment was slightly different from the one used in the fMRI study: Each critical trial only contained the same word presented twice in a row either in the same modality or in different modality. Here, the time-course of neural activity was extracted for each word and the RSE was computed by contrasting the neural responses obtained on two consecutive stimuli.

The protocol contained four conditions of same-word pairs that were randomly presented among the fillers. The four conditions reflected the modality of the word that was repeated twice in a row (AA: both presentations were in the auditory modality, VV: both presentations were in the visual modality, AV: the first presentation was in the auditory modality and the second was in the visual modality, VA: the first presentation was in the visual modality and the second was in the auditory modality), as illustrated in **Fig. 1B**. Four types of RSE were estimated by contrasting the brain activity measured during the first and the second presentation of the word. The task was presented in five blocks that lasted ∼3.8 min each. Each block contained 10 critical trials (pair of words) from each condition. They were randomly presented among 80 fillers (10 per type of fillers) and 16 catch trials (eight ###### and eight beep sounds), with a constraint that no condition was presented twice in a row. The duration of the written inputs was 550 ms while the duration of the spoken inputs depended on the file duration (ranged from 307 ms to 555 ms). The interval between two stimuli was jittered between 500 ms and 600 ms. During the task, the participants were not informed about the existence of word pairs and were requested to detect the catch trials by pressing the response button. Note that, to reduce fatigue and cognitive demands, the task applied on epileptic patients was simpler than the one applied in the fMRI study conducted on healthy participants.

#### sEEG Data Acquisition

Intracerebral electrodes were implanted for clinical purposes, using 13 to 17 depth-electrodes, each containing 10 to 15 recording sites of 2 mm in length and separated by a 1.5 mm distance (0.8 mm in diameter; Alcis, Besançon, France). Patients were laying on their hospital bed with a laptop in front of them (approx 80 cm). Data were recorded using a 256-channels Natus amplifier (Deltamed system) and sampled at 1000 Hz.

#### sEEG Electrode Localization and Selection

To precisely localize the channels, a procedure similar to the one used in the iELVis toolbox was applied (81). First, we manually identified the location of each channel centroid on the post-implant CT scan using the Gardel software (82). Second, we performed volumetric segmentation and cortical reconstruction on the pre-implant MRI with the Freesurfer image analysis suite (http://surfer.nmr.mgh.harvard.edu/). Third, the post-implant CT scan was coregistered to the pre-implant MRI via a rigid affine transformation and the pre-implant MRI was registered to the MNI template (MNI 152 Linear), via a linear and a non-linear transformation from SPM12 methods, through the FieldTrip toolbox (83). To identify sEEG contacts that are within the left-vOT, a bounding box was created to cover the left-vOT by considering individual variability in this functional area (84). Specifically, the individual ROIs obtained in the fMRI visual localizer were used to define the boundaries of the bounding box, where X E [-64.2, -25.4], Y E [-72.1, -21.0] and Z E [-35.8, -4.0] in MNI space. The sEEG electrode contacts located within the bounding box were further selected based on significant high-frequency activity to visual words to confirm the corresponding visual word processing.

#### sEEG Data Analysis

The pre-processing of sEEG data was conducted using Brainstorm (85). For each patient, power spectral density (PSD) was estimated using Welch’s method and visually inspected by two authors (SW and ASD) to mark outliers in PSD as noisy electrode contacts. None of the left-vOT electrode contacts were rejected. Notch filters were applied at 50, 100, 150, 200 and 250 Hz. The signals were then high-pass filtered with 0.3 Hz using a Kaiser-window linear phase FIR filter of order 12376. The monopolar and bipolar data were visually inspected for marking bad segments by two authors (WS and ASD) who were blinded to trial labels. The signals were then segmented into epochs of -300 ms to 600 ms locked to stimulus onset. These epochs were imported in Multi-patient Intracerebral data Analysis (MIA) toolbox (51) for estimating high-frequency activity (HFA; 70-150 Hz with steps of 10Hz) to conduct multi-patient permutation tests. All analyses were conducted using a bipolar montage with a 7 cycles Morlet wavelet and a baseline from 300 ms to 10 ms before trial onset to exclude edge effects. The multi-patient permutation tests were conducted for each type of RSE with 1000 iterations and the significant threshold *p* < .05 (50, 51, 86; SI Appendix).

## Acknowledgments

This work was supported by the French Ministry of Research: ANR-13-JSH2-0002 and ANR-19-CE28-0001-01 (to C.P.), ANR-16-CONV-0002 (ILCB), ANR-11-LABX-0036 (BLRI) and the Excellence Initiative of Aix-Marseille University (A*MIDEX). We thank Dr. Kenneth Pugh for his helpful suggestions regarding the protocol design. This work was performed in the Center IRM-INT (UMR 7289, AMU-CNRS), platform member of France Life Imaging network (grant ANR-11-INBS-0006). Centre de Calcul Intensif d’Aix-Marseille is acknowledged for granting access to its high-performance computing resources.

## Author Contributions

C.P, S.W, A-S.D, J-L.A., and A.T designed research; S.W., A-S.D., V.C., C.P, J.S, and B.N performed research; S.W., A-S.D. and M.M. analyzed data, S.W. and C.P. wrote the original draft, S.W., C.P., A-S.D, V.C., M.M, and J.S reviewed and edited the original draft, C.P. acquired funding.

## Competing Interest Statement

The authors declare no competing interest.

## Classification

Major: Biological Sciences. Minor: Psychological and Cognitive Sciences

## Notes

### Competing Interest Statement

The authors have declared no competing interest.

