## Supporting Information for "Revealing the co-existence of written and spoken language coding neural populations in the left-ventral occipitotemporal cortex"

Address: Laboratoire Parole et Langage (UMR 7309). 5 Av. Pasteur, 13100 Aix-en-Provence, France

Phone number: +33 601323435

**This PDF file includes:**

Supporting Information Text

Figures S1 to S5

SI References

### Supporting Information Text

#### Supplementary Results

**Identification of individual ROI within the left-vOT ($ROI_{INDvOT}$).** In each participant, the peak coordinates and the corresponding T-value were obtained from the first-level (individual) comparison of *words* and *consonant strings*. Participants whose peak activation was significant at $p < 0.001$ ($T > 3.32$) were included, resulting in 18 participants with peaks in the ranges of MNI x = [-56, -33], y = [-64, -29], z = [-28, -12] (**Fig. S1 A**). The individual ROIs were defined by creating an 8 mm sphere (389 voxels) centered at the peak coordinates. The 8mm radius resulted in a spherical ROI with a similar volume as $ROI_{GRPvOT}$ which contained 311 voxels. As shown in **Fig. S1 B**, the individual ROIs ($ROI_{INDvOT}$) had higher activation in *words* than in *consonant strings* ($p < 0.0007$).

**Examination of individual ROIs responses to spoken inputs and its repetition suppression pattern**. The ROI-based analysis conducted on the $ROI_{INDvOT}$ confirmed the results that we obtained on the $ROI_{GRPvOT}$. As shown in **Fig. S1 C**, in the auditory task, $ROI_{INDvOT}$ showed higher activation in response to *spoken words* and *spoken pseudowords* than to s*crambled stimuli* (all *p*s < 0.0023 unc.) and no difference between *spoken words* and *spoken pseudowords* was found ($p > 0.39$ unc.). In the repetition suppression protocol, $ROI_{INDvOT}$ showed significant within-modal RSEs in both visual (*SameVV - DiffVV*) and auditory (*SameAA-DiffAA)* modalities ($p < 0.016$ unc. and $p < 0.046$ unc., respectively; **Fig. S1 D**). As in the main analysis, no cross-modal RSE was observed (all *p*s > 0.09 unc.).

**Voxel-wise univariate analysis conducted in the auditory task**. The whole-brain analysis revealed strong activation in the left-vOT for both the *spoken pseudowords - scrambled stimuli* and *spoken words - scrambled stimuli* contrasts (FWE $p < 0.05$, voxel-wise $p < 0.005$; **Fig. S2** axial views), whereas no significant activation was found for the *spoken words - spoken pseudowords* contrast. The bilateral temporal, precentral and postcentral regions also showed the same activation pattern (FWE p < 0.05, voxel-wise $p < 0.005$; **Fig. S2** surface views).

**Validation of the repetition suppression protocol**. In our main analyses, the examination of the RSEs within the left-vOT only revealed significant within-modal RSEs (**Fig. 3**). Here, we validated the existence of the pure auditory RSE and cross-modal RSEs by extending the ROI-based analysis to left temporal regions. First, we conducted the analysis on the left and right STG representing the primary auditory cortex. Both STGs were extracted from the AAL template (1) and confined with the group-averaged gray matter mask. As expected, both left and right STG only showed auditory RSE (**Fig. S3 A** and **B**; $p < 0.0039$ and $p < 0.0052$, respectively; permutation tests with FWE correction for each ROI). Second, we identified a focus in the left pSTS that is considered as high-level multimodal audiovisual integration area, according to a meta-analysis study (2). This ROI was defined by creating an 8 mm sphere centered at the focus' coordinates (MNI -54, -44, 8). It also overlapped with the foci involved in the integration of speech and orthographic information reported by van Atteveldt et al. (3) (TAL -54, -48, 9) and Rueckl et al. (4) (MNI -48 -42 6). As shown in **Fig. S3 C**, this ROI showed significant within-modal and cross-modal RSEs (all *p*s < 0.045; permutation tests with FWE correction).

**MVPA results using Linear Discriminant Analysis (LDA) and Gradient Boosting Classifier (GBC).** To confirm the MVPA results using linear and non-linear SVMs, the searchlight analysis was conducted using another simple linear classifier Linear Discriminant Analysis (LDA) and a non-linear tree-based Gradient Boosting Classifier (GBC), which has a very flexible decision boundary. In line with the results of SVMs, the linear classifier LDA only showed two significant clusters with above-chance-level accuracies for written inputs (FWE $p < 0.05$, voxel-wise $p < 0.005$; **Fig. S4 A**). The first cluster was centered in the posterior fusiform gyrus extending into the inferior occipital cortex (peak MNI -40, -76, -12). The second one was centered in the anterior fusiform gyrus and largely overlapped with the left-vOT (peak MNI -47, -40, -22). The non-linear GBC revealed a significant cluster that had above-chance-level accuracy for spoken inputs around the left-vOT (**Fig. S4 B** and **4C**; FWE $p < 0.05$, voxel-wise $p < 0.005$; peak MNI -39, -20, -22), as well as three clusters with above-chance-level accuracies for decoding written inputs. These three clusters are at the similar locations as those obtained in the linear SVM (FWE $p < 0.05$, voxel-wise $p < 0.005$; peak MNI -49, -64, -7; -49, -43, -26 and -35, -85, -21; **Fig. S4 B** and **4C**). Note that neither linear LDA nor non-linear GBC led to an above-chance-level accuracy in the cross-modal conditions.

**Multimodal regions revealed by MVPA decoding of stimulus lexicality.** To search for multimodal clusters in the high-order language regions, we created a mask covering the left middle and superior temporal gyri, left supramarginal gyrus and left angular gyrus from the AAL template (1). The mask was then confined by the group-averaged gray matter mask. The searchlight accuracy maps were first estimated for each of the four decoding conditions with FWE $p < 0.05$, voxel-wise $p < 0.005$ (**Fig. S5 A**). Then, we identified the areas that showed a significant decoding performance in all conditions by using a conjunction analysis (**Fig. S5 B**). The result revealed two clusters centered at the left pSTS (MNI -61, -40, 1) and left temporoparietal junction (MNI -48, -70, 19), which are involved in multimodal language processing (5–7). As shown in **Fig. S5 C**, the cluster in the left pSTS revealed by searchlight MVPA overlapped with ROI defined from the meta-analysis by Erickson et al. (2) (**Fig. S3 C**).

### Supplementary Methods

***fMRI Data Pre-processing.*** The T1-weighted image was corrected for intensity non-uniformity with N4BiasFieldCorrection in ANTs (8, 9), and used as T1w-reference throughout the workflow. The T1w-reference was then skull-stripped. The brain-extracted T1w was used for segmentation of cerebrospinal fluid (CSF), white-matter (WM) and gray-matter (GM) using fast (FSL 5.0.9). Volume-based spatial normalization to the standard MNI space was performed through nonlinear registration with antsRegistration, using brain-extracted versions of both T1w reference and the T1w template (MNI152NLin2009cAsym). For functional images, the fieldmap distortion correction was performed based on a phase-difference map. The functional images were then co-registered to the T1w reference using flirt (FSL 5.0.9) with the boundary-based registration (10) with nine degrees of freedom. Head-motion parameters were estimated before any spatiotemporal filtering using mcflirt (FSL 5.0.9). Fieldmap distortion correction, head-motion correction, BOLD-to-T1w co-registration, and spatial normalization were carried out in a single interpolation step by composing all the pertinent transformations. The pre-processed BOLD data were then used to calculate several confounding time series, including framewise displacement (FD), the mean signals within the white matter and the CSF, and a set of principal components of white matter and CSF that were extracted by the aCompCor method (11).

***Multi-patient permutation tests (sEEG).*** For each trial, time-frequency power was computed on consecutive 10 Hz bands between 70 and 150 Hz with a 7 cycles Morlet wavelet. Baseline correction was applied at each 10 Hz band by calculating a z-score relative to activity during the baseline from 300 ms to 10 ms before trial onset to exclude edge effects. The statistical tests were conducted on the z-score data and were corrected for multiple comparisons in the time domain by estimating a minimum duration threshold for consecutive significant t-values. Specifically, for between-condition tests, the trials of two conditions were randomly permuted at each time point for calculating the maximum number of consecutive time points passing the significance threshold ($p < .05$ unc.). The permutations were conducted 1000 times to obtain a distribution of 1000 maximum numbers. The minimum duration threshold was defined at the right-tail 95% quantile of this distribution.

**Supporting Figures S1 to S5**

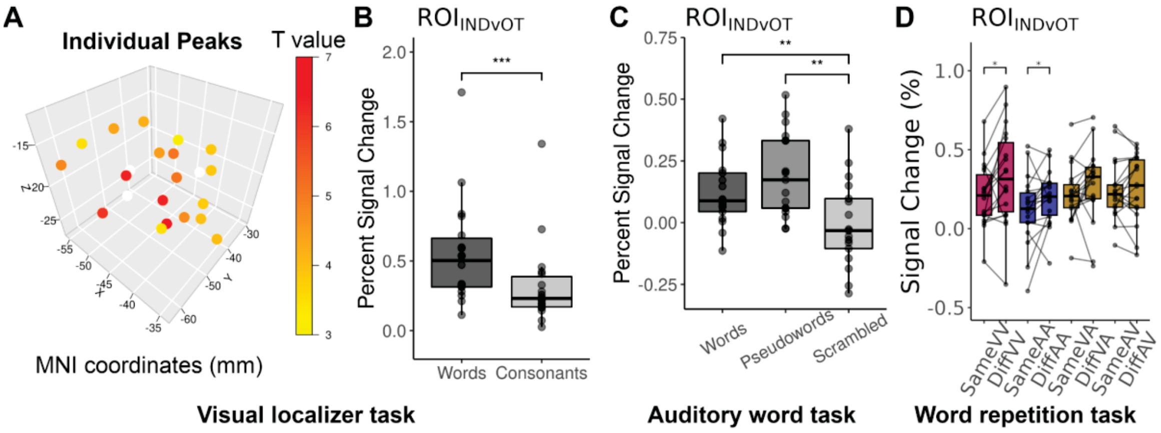


**Fig. S1.** Identification of individual ROIs within the left-vOT (ROI$_{INDvOT}$) and its functional profiles. (A) 3D scatters showing the individual peaks that were significant at $p < 0.001$. In each participant, the individual ROI was created as a sphere centered at the peak coordinates with a radius of 8mm. (B) The ROI$_{INDvOT}$ showed higher activation to *words* compared to *consonant strings*. (C) The ROI$_{INDvOT}$ showed higher activation to *spoken words* and *spoken pseudowords* compared to s*crambled stimuli*. (D) The ROI$_{INDvOT}$ showed significant within-modal visual and within-modal auditory RSEs while no cross-modal RSE was observed.

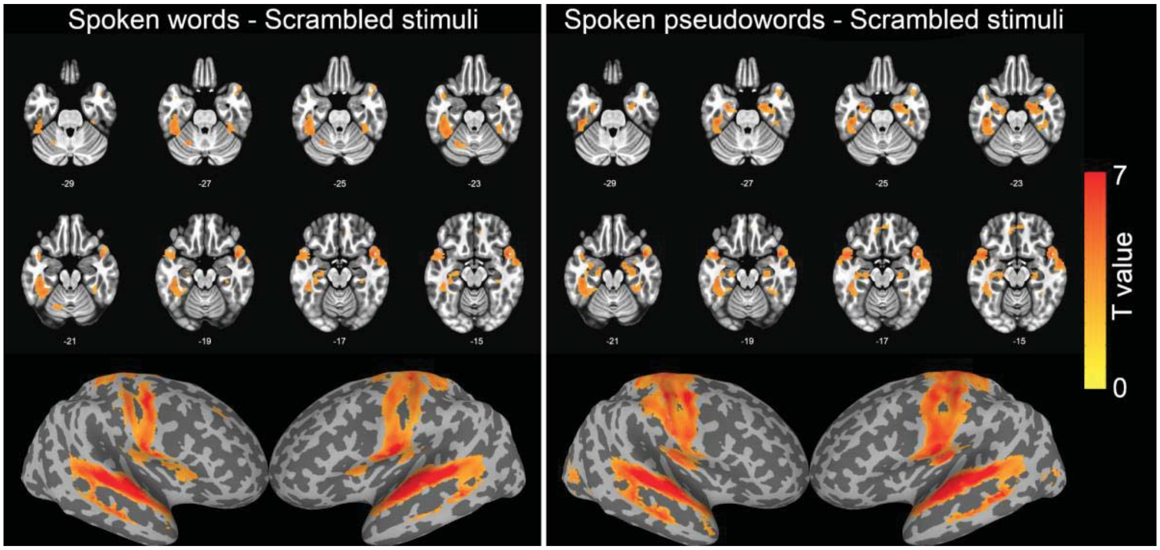


**Fig. S2.** Activation maps of brain regions that showed significant activation in the *spoken pseudowords* - *scrambled stimuli* and *spoken words* - *scrambled stimuli* contrasts, including left-vOT, bilateral temporal, precentral and postcentral regions (FWE $p < 0.05$, voxel-wise $p < 0.005$).

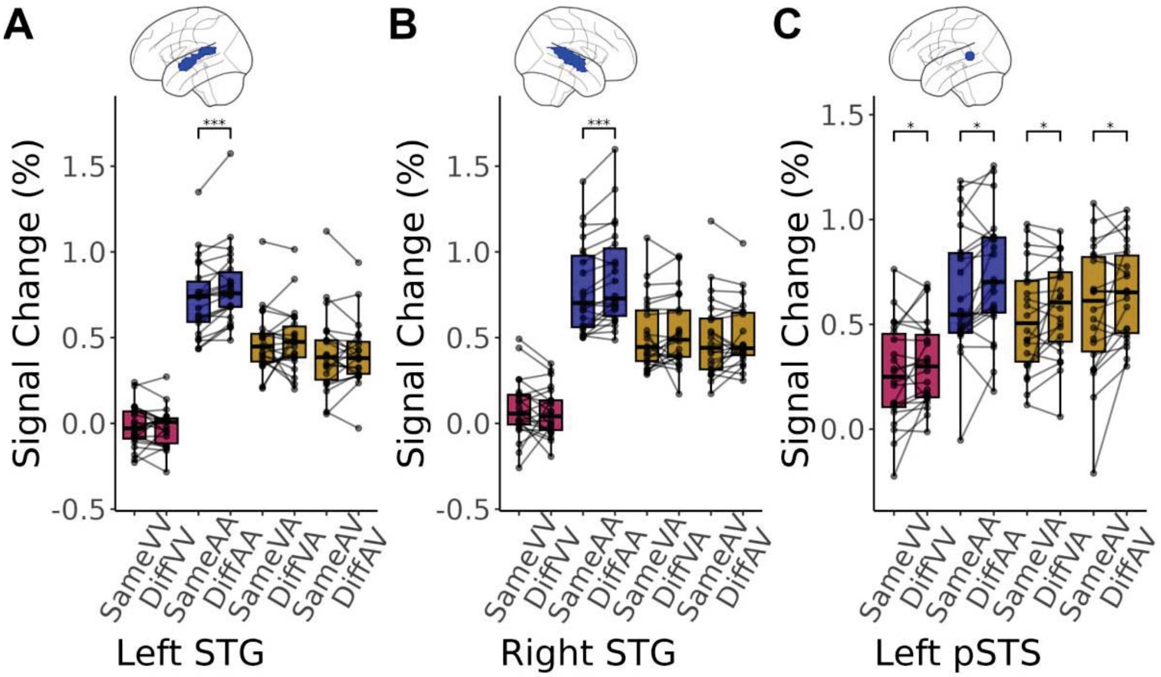


**Fig. S3.** Repetition suppression effects in primary auditory cortex (bilateral STGs in AAL template) and a multimodal language region in left pSTS (2). (A) and (B) Both left and right STGs only showed significant within-modal auditory RSE. (C) The ROI in left pSTS showed both within- and cross-modal RSEs. *: $p < 0.05$; **: $p < 0.01$; ***: $p < 0.005$.

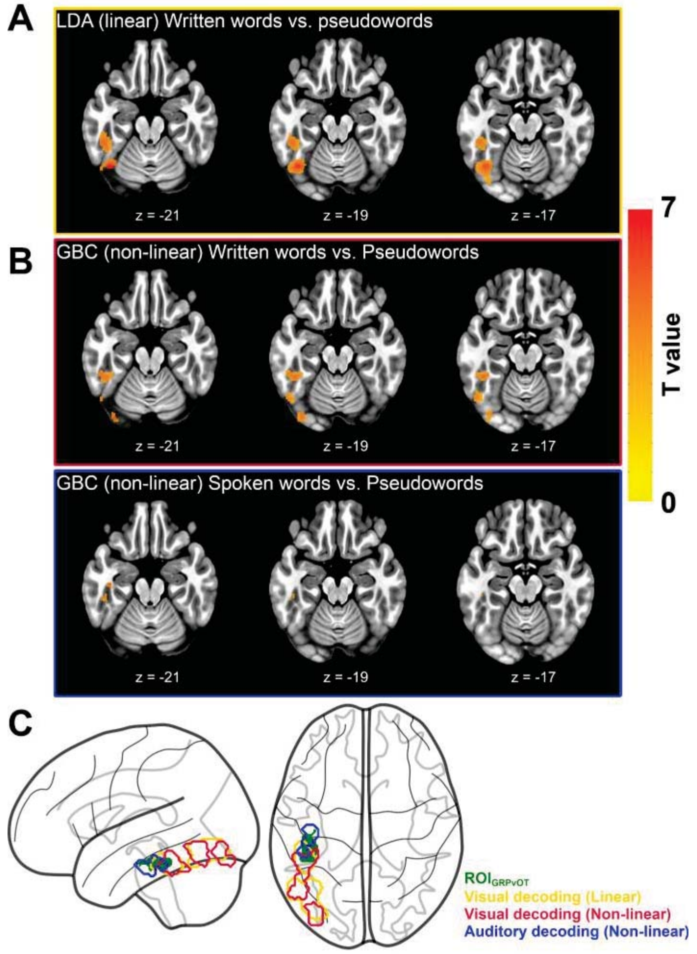


**Fig. S4.** Results of searchlight MVPA for lexicality decoding in the left-vOT (A) The linear classifier LDA revealed two significant clusters that have above-chance-level accuracies for written inputs (FWE $p < 0.05$, voxel-wise $p < 0.005$). (B) The non-linear classifier GBC revealed three significant clusters for written inputs and one significant cluster for spoken inputs (FWE $p < 0.05$, voxel-wise $p < 0.005$). (C) Glass brain showing the overlap between the $ROI_{GRPvOT}$ (green patch) and the clusters that represented above-chance visual (yellow and red contour for LDA and GBC, respectively) and auditory decoding performance (blue contour).

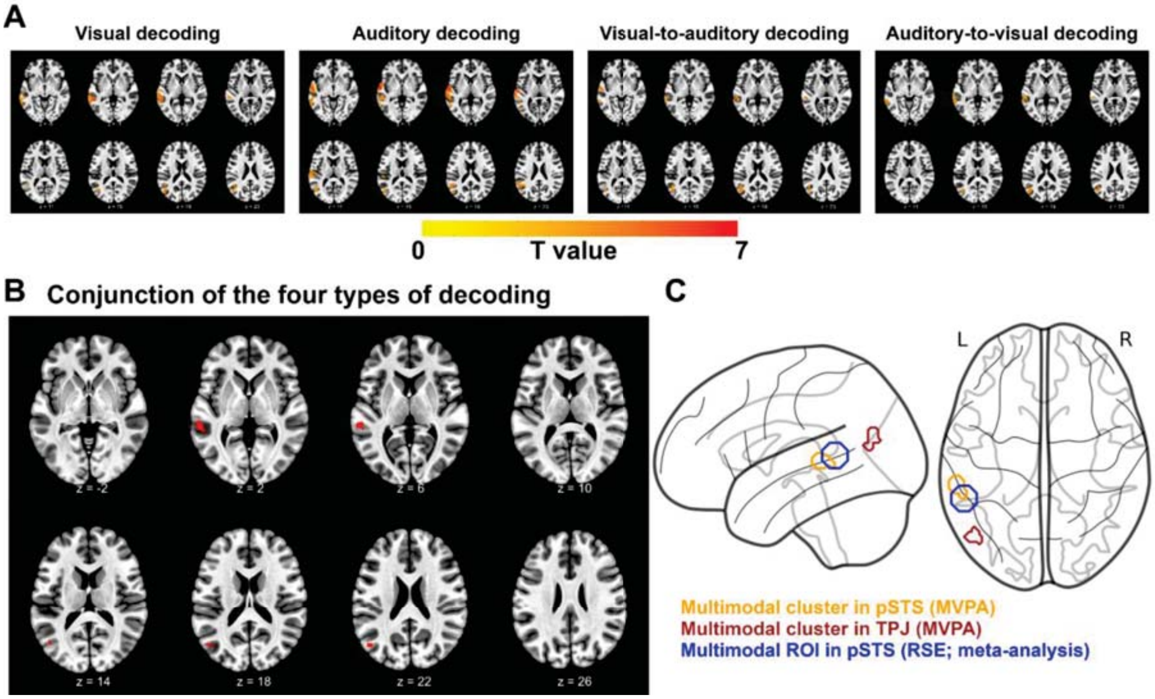


**Fig. S5.** Searchlight accuracy maps. (A) Brain regions that showed above-chance level accuracy in visual decoding, auditory decoding, visual-to-auditory decoding and auditory-to-visual decoding (FWE $p < 0.05$, voxel-wise $p < 0.005$). (B) Two clusters in the left pSTS and TPJ that showed above-chance level accuracies in both within- and cross-modal decoding conditions. (C) The multimodal cluster in the left pSTS (orange contour) and the left TPJ (red contour) revealed by MVPA. The former region overlapped with the multimodal ROI that showed both within- and cross-modal RSEs (blue contour).
